# Selective manipulation of the host RNA-polymerase transcription fidelity increases phenotypic mutations in an insect *Parvovirus*

**DOI:** 10.1101/2024.01.15.575645

**Authors:** Thomas Labadie, Guillaume Cambray

## Abstract

In the dynamic dance of evolution, organisms are often faced with fluctuating environments to which adaptation through selection of traditional heritable genetic mutations can be limiting. In this study, we unveil a refined mechanism of non-heritable variability in a virus with a compact DNA genome. We discovered that the genome of the *Junona cœnia* densovirus (JcDV) experiences a 10-fold lower transcription fidelity than the rest of the host’s transcriptome, despite the shared transcription machinery. We found that the virus’ capsid proteins interact with the host’s RNA Polymerase II, and further show that truncating these proteins through early stop codons largely restore the transcriptional fidelity of viral genes. These observations suggest a potential mechanism for the selective manipulation of transcription accuracy. We also pinpoint specific sequence contexts that may provide other knobs to finely control the local transcription fidelity. We estimate that this lower transcriptional fidelity results in more than 7% of the viral proteome bearing at least one non-synonymous mutation. The production of non-heritable functional diversity by hijacking the transcriptional machinery might be a refined strategy to enhance short term adaptation within the complex and ever-changing host-parasite interface, and might be shared by other genetic parasites. Our findings shed light on a virus-specific transcription fidelity control mechanism, expanding our understanding of adaptive strategies in biological entities.

## INTRODUCTION

Living organisms are torn between the need to maintain their genetic integrity, which enabled their survival, and the necessity of producing the variations required for adaptation to changing environments. Mistakes made during transcription and subsequent translation can lead to the production of truncated, misfolded or dysfunctional proteins that will modify or alter the link between genotype and phenotype. Unlike their genetic counterparts, these variations–collegially referred to as phenotypic mutations–are non-inheritable. The most studied source of phenotypic mutations is translational errors, using model reporter systems (1–3) such as programmed frameshifting or stop codon read-through, that give signals without need for mass spectrometry methods which are still limiting. Advances in RNA sequencing recently unveiled RNA mutation rates, which are higher than DNA mutation rates, as an important source of phenotypic mutations (1, 4–6). Yet the mechanism providing these mutations are not well understood and has mainly been studied in the context of artificial, error-prone transcription systems (1–3).

Despite their sporadic appearance, such mutations can elicit increased phenotypic variance around a constant genotype (**7**). They are suspected to confer selective benefits for the acquisition of complex traits requiring multiple genetic mutations (**8**) such as increased stability (**9**). In addition, phenotypic mutations are predicted to exert a stronger impact on the evolution of organisms with large population size (**10**) such as viruses, thereby enabling alternative mechanisms for virus adaptation to constant changes of hosts and environments (4). Typically, DNA viruses hijack the host DNA-dependent RNA polymerase II (RNAP-II) in eukaryotes to produce messenger RNA (mRNA). This provides an opportunity to compare transcriptional mutations in the context of the host and parasite interaction, who both have distinct evolutionary rate and population sizes but share the same transcription machinery.

In this study, we explored the transcription fidelity of the *Junonia coenia* Densovirus (JcDV), a small insect *Parvovirus*, which infects larvae from the fall armyworm, a major pest crop worldwide. This virus holds great potential for controlling populations of nefarious insects, although we are lacking refined molecular understanding to firmly establish safety and control for such applications. It has a small (6,048 nucleotides), single stranded DNA genome (**Figure 1A**), which yields two ambisense mRNAs (**11**) during transcription. One mRNA encodes four structural protein isoforms (VP1-4) that define the viral capsid (**12**). The second mRNA encodes three non-structural proteins, with NS1 being clearly involved in genome replication through its helicase and nickase activities (**13**) and NS2-3, whose role remains to be elucidated (**14**). In addition, the viral genome harbours inverted-terminal repeat sequences at both termini, implicated in replication and packaging processes (**15, 16**). Our study revealed that viral genes are transcribed with significantly higher error rates than cellular genes. We were able to demonstrate that this higher error rate during viral gene transcription is associated with the expression of viral capsid proteins (VP), potentially facilitated by an interaction between the capsid proteins and the RNAP-II interaction.

**Figure 1.**
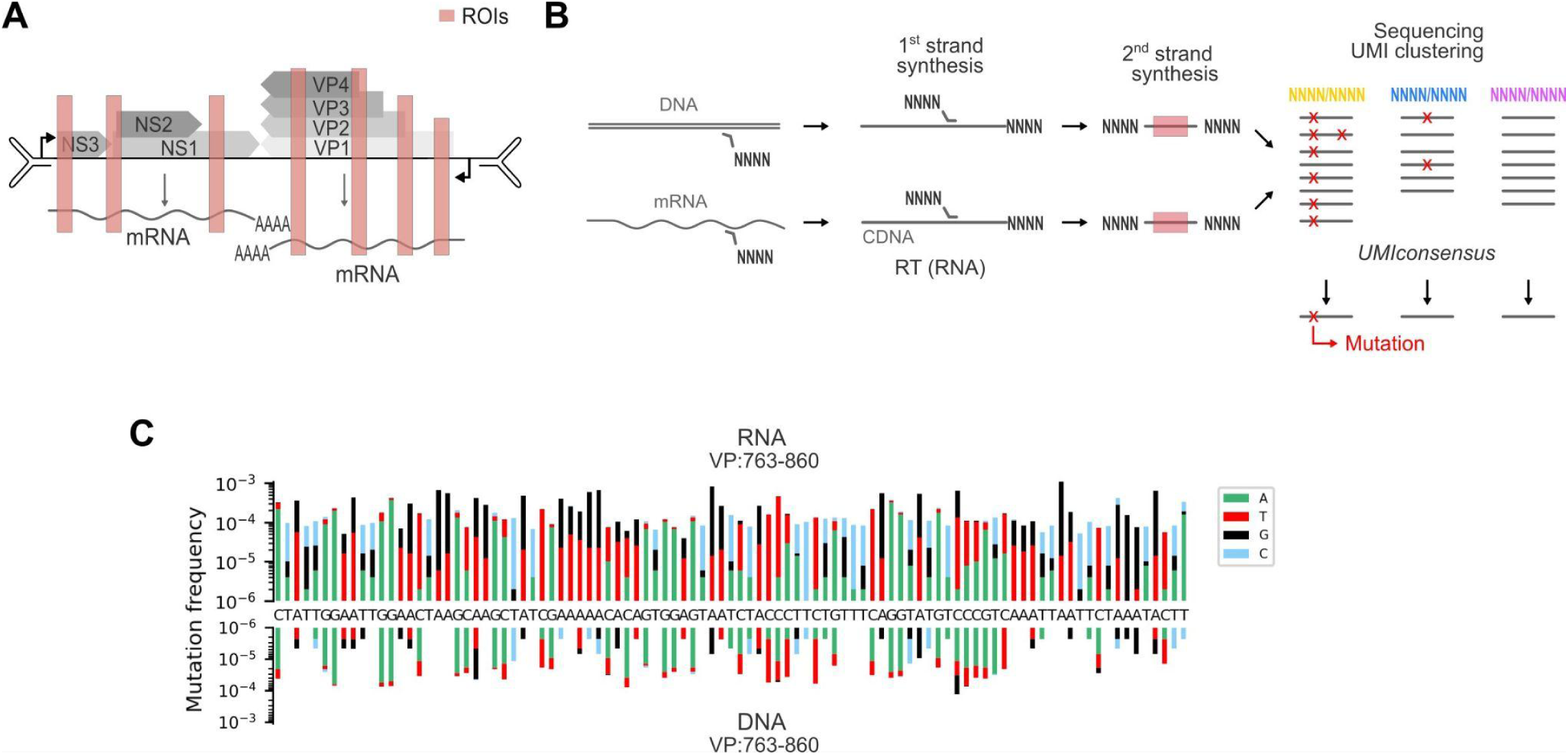
Measure of transcription fidelity. (A) Diagram of the JcDV genome organisation and associated transcripts. Region of interest (ROI) analysed by targeted deep sequencing are highlighted in red. (B) UMI (NNNN) were fused to ROI targeting primers used for reverse transcription (RT) of mRNA or first and second second strand synthesis (DNA and cDNA). After sequencing, UMI clustering allows us to retrieve all reads copied from single RNA or DNA molecules, which provides a stringent consensus sequence for each UMI cluster. (C) Frequency of substitutions/nucleotide measured at all positions in mRNA (top) or DNA (bottom) for the ROI VP:763-860. Reference nucleotides are indicated at each position (middle).

## MATERIAL AND METHODS

### Cell line, infection and transfection

The insect cell line Ld652Y (17) was maintained at 28°C in TC-100 medium (Gibco) supplemented with 10% heat-inactivated fetal calf serum (FCS, Thermo Scientific) and an antibiotic/antimycotic cocktail (10 U/ml penicillin, 10 μg/ml streptomycin, and 25 fg/ml amphotericin B; Gibco). Ld652Y cells were originally derived from *Lymantria dispar* ovaries and are most sensitive to JcDV infection (18). A JcDV virus stock was produced by infecting Ld652Y cells at a multiplicity of infection (MOI) of 1, and harvesting the cell supernatant at 5 to 7 days post infection, once all cells were lysed. Cell debris were removed by centrifugation (500 x g for 5 minutes), and the virus stock was aliquoted and stored at −80°C.

Ld652Y infection experiments were all performed using the same virus stock. Briefly, flask-grown cells were harvested, and seeded in new T75 cm^2^. The next day, when cell growth reached ~70% confluency, the cell culture media was removed, and cells were infected with a 3 mL inoculum of JcDV stock diluted in TC-100 to effect a MOI of 10. After 30 minutes incubation at room temperature, the inoculum was removed and replaced by 10 mL of TC-100 supplemented with 10% FCS and antibiotic/antimycotic cocktail. Cells were then incubated at 28°C.

For transfection, Ld652Y were first seeded in T25 cm^2^ flasks. The next day, cell culture medium was removed and replaced by fresh TC-100 supplemented with 10% FCS and antibiotic/antimycotic cocktail. Cells were then transfected with plasmid DNA (4µg per plasmid per flask) using FuGENE HD (1:3 ratio; Promega) and cultured at 28°C.

### Plasmids

The WT JcDV clone corresponds to the published *Junonia coenia densovirus* isolate Oxford (15) cloned into a pIZ vector (Zeocin resistance), along with a rapporter fluorescent mScarlet gene under the control of a *OpIE*2 promoter. Compared with the published sequence, a BsaI digestion site contained in the NS3 gene was removed by inserting the synonymous substitution AGG to CGT at position 1007 of the genome using site directed mutagenesis. This vector was used for initial transfection of Ld652Y cells and virus stock preparation.

To introduce early stop codons in viral genes, a donor vector containing a JcDV genome without terminal hairpin sequences (allowing whole genome amplification by PCR) encompassed by two SapI digestion sites, was mutated by site-directed mutagenesis (Q5 Site-Directed Mutagenesis Kit, New England Biolabs, E0554). To assemble full JcDV genomes after site directed mutagenesis, an acceptor plZ vector was constructed, containing the missing hairpin terminal sequences encompassing two SapI digestion sites, a zeocin resistance gene Sh ble, a mScarlet reporter gene under the control of an OpIE2 promoter. Plasmids bearing mutant JcDV genomes were obtained by subcloning the mutated donor insert into the acceptor vector, using a golden gate assembly strategy (19) and a type-II restriction SapI enzyme (New England Biolabs, R0569). A similar cloning strategy was used to build vectors bearing a JcDV viral genomes with fragments of cellular genes (~160 nucleotides) fused in the VP genes. This time, a a different plZ acceptor vector was built, with a complete JcDV, except for a 110 base pair window located in position 746 of the VP gene, replaced by two BsaI digestion sites. The mScarlet reporter gene was also replaced by an additional VP gene for trans complementation. Cellular genes were amplified by PCR (Q5 High-Fidelity DNA Polymerase, New England Biolabs) after DNA extraction (Quick-DNA Miniprep Kit, Zymo Research) from the LD652Y cells using primers bearing BsaI restriction sites in their 5’ tail, and cloned into the accepting vector using a golden gate cloning strategy (20).

### Library preparation and sequencing

Oligonucleotides were synthesised by Integrated DNA technologies or Eurofins genomics.

Infected or non-infected Ld652Y cells were resuspended in PBS buffer at 24 hours (unless specified) post infection or transfection, after two PBS washes of the cell monolayers. Cells were pelleted by centrifugation (500 x g for 5 minutes) and lysed for RNA and DNA extraction using the Quick-DNA/RNA Miniprep Plus kit (Zymo Research, D7003). RNA and DNA were eluted in water, followed by DNaseI treatment of the RNA for 10 min at 37°C (New England Biolabs, M0303), and removal of the DNaseI using the RNA clean up kit (New England Biolabs, T2040).

For RNA sequencing, a reverse transcription step was performed using 15 µg RNA input and the superscript IV reverse transcriptase (Thermo Scientific) with a 2 µM reverse gene specific primers containing an additional UMI sequence (8 random nucleotides) and partial nextera adaptor sequences in their 5’ tails. Second strand synthesis was performed using the entire reverse transcription pot as input into a one cycle PCR reaction (98°C: 0’30 sec, 65°C: 0’15 sec, 72°C: 1’0 min) with a Q5 high fidelity DNA polymerase (New England Biolabs, M0491) and 10 µM forward gene specific primers also containing an additional UMI sequence (8 random nucleotides) and partial nextera adaptor sequences in their 5’ tails.

For DNA sequencing, targeted DNA barcoding with UMI was performed using a two cycles PCR reaction (98°C: 0’30 sec; then two cycles of : [98°C: 0’10 sec, 65°C: 0’15 sec, 72°C: 0’15 sec]; and a final 72°C: 1’0 min) with 1 µg DNA input, a Q5 high fidelity DNA polymerase (New England Biolabs, M0491), and 10 µM forward and reverse gene specific primers containing an additional UMI sequence (8 random nucleotides) and partial nextera adaptor sequences in their 5’ tails.

For both RNA and DNA barcoded with UMIs, non incorporated primers were removed using the Monarch PCR and DNA clean up kit (New England Biolabs, T1030), using a binding buffer ratio to DNA of 3:1. Then NGS libraries were then amplified by PCR using a Q5 high fidelity DNA polymerase, 100 µM indexed P5 and P7 adapters, and total purified barcoded DNA as input. Various PCR cycles (**Supplementary Figure S1A**) of 98°C: 0’30 sec; then cycles of : [98°C: 0’10 sec, 72°C: 0’30 sec]; and a final 72°C: 3’0 min, were required depending on the input template.

Final library purification was performed using the ProNex Size-Selective DNA Purification System (Promega, NG2001) and eluted in water before libraries pooling and sequencing using a paired-end 150 method on a Illumina NovaSeq System by Novogene.

### Bioinformatic analysis

We built a python package called UMIconsensus to process paired-end sequencing reads into analysable mutation profiles. The pipeline essentially takes 3 main inputs: demultiplexed fastq files containing the sequencing reads for a particular ROI, the two sequences flanking the ROI up to the introduced UMIs and the sequence of the ROI itself. The processing pipeline starts by calling NGmerge to merge paired overlapping reads into single sequences (21). NGmerge uses an experimentally informed model to compute base qualities that compound information from aligned paired reads, and therefore provide a first layer of error correction at overlapping positions. Paired reads that do not overlap or whose overlapping region comprises too high a mismatch fraction are rejected. Successfully merged reads are then processed using CutAdapt (22) to identify the two flanking sequences and extract the sequence of the ROI and its associated UMIs. Sequences that are not properly resolved by CutAdapt or whose extracted UMIs do not match the expected sizes are rejected. Global pairwise alignments (Needleman-Wunsch) of the identified ROI sequences and qualities to the reference ROI sequence are then produced using parasail (23). Once so processed, reads are grouped on the basis of their concatenated UMIs. It has been described that sets of UMIs identified in any sequencing run are typically enriched in sequences that are too close from one another and likely correspond to single original UMI sequences that have been diversified by amplification and sequencing errors occuring after their physical introduction in the sequenced molecules. A number of algorithms to cluster related UMI have been proposed in UMItools (24) and reimplemented in the faster UMIcollapse (25). Within a given UMI cluster, the previously generated pairwise alignments are consolidated in a multiple alignment data structure that is used to dynamically call a consensus sequence for that molecule according to two user-specified thresholds. The first threshold defines the minimal quality below which individual bases in a sequencing read are excluded from analysis. The second threshold defines the minimal number of quality-filtered bases acceptable to call a consensus at a given position. The consensus is defined as the most represented base at a position. In case of ties, the reference base at this position is given precedence and then the base associated with highest average quality. This procedure permits to discriminate PCR-based mutations that arose after the introduction of the UMIs and sequencing errors from true mutations present in the original nucleic acid molecule with a stringency that depends on the chosen clustering method and filtering thresholds. Once reconstructed, the UMI-based consensus sequences are finally used to construct a large multiple alignment from which the frequencies of mutations can be easily computed and plotted. All the data presented in this work are produced using the ‘adjacency’’ algorithm of UMIcollapse, a quality threshold of 35 and a coverage threshold of 3. Variation of these parameters have little impact on the conclusion of this study, as detailed in **Supplementary Figure S1** and **Supplementary Table S1**.

### Quantification of transcript abundance by quantitative PCR (qPCR)

Extraction of the DNA and RNA were performed using a Quick-DNA/RNA Miniprep Plus kit (Zymo Research, D7003), followed by DNaseI treatment of the RNA for 10 min at 37°C (New England Biolabs, M0303), and removal of the DNaseI using the RNA clean up kit (New England Biolabs, T2040). For the RNA quantification, a reverse transcription step prior qPCR was performed using the superscript IV reverse transcriptase (Thermo Scientific) with random hexamers. The cDNAs obtained were then quantified with a real time qPCR, along with extracted DNA, using the Luna universal qPCR Master Mix (New England Biolabs, M3003) for cellular genes or the Luna universal probe qPCR Master Mix (New England Biolabs, M3004) for the viral transcripts. For absolute quantification of targeted genes, a standard curve was produced by amplifying a 10-fold serial dilution of plasmids containing the target sequences at known concentrations. Quantification values for RNA transcripts were then normalised by quantification values obtained for their respective DNA encoding genes.

### Immunoprecipitation assays

Cell monolayers were washed twice with a PBS solution and lysed for 10 min at 4°C with an immunoprecipitation lysis buffer (25 mM Tris, 150 mM NaCl, 1 mM EDTA, 5% v:v glycerol, 1% v:v NP40 /IGEPAL GA-630, pH7.4 adjusted with HCl) containing 1X Protease Inhibitor Cocktail (Sigma-Aldrich 11873580001). Cell lysates were then centrifuged at 13,000 × g for 10 minutes at 4°C to pellet the cell debris. The soluble extracts were incubated on a roller with anti-VP, NS1, NS2 or anti-RNAP-II (CTD Antibody, clone 8WG16, Sigma-Aldrich 05-952-I) antibodies overnight at 4°C, followed by protein G coupled magnetic beads at room temperature for 10 minutes (PureProteome Protein G Magnetic Bead System, Millipore LSKMAGG). The beads were removed using a magnetic rack, and were further washed four times with a PBS buffer containing 0.1% Tween20. After the final wash, equal amounts of PBS and 2X Laemmli sample buffer were added to the beads and heated for 5 min at 95°C. Samples were then proceeded for western blot analysis.

### Western blot assays

After cell lysis using an immunoprecipitation buffer (see section immunoprecipitation assays), or after immunoprecipitation assays, samples were diluted in 4X Laemmli Sample Buffer (Thermofisher scientific) to a final 1x concentration and incubated at 95°C for 5 min. Proteins were then separated on a 4-12% SDS gel and blotted on a nitrocellulose membrane. When necessary, the membranes were cut horizontally for incubation with different antibodies. The primary rabbit anti-VP, anti NS1, anti NS2 antibodies (a gift from Anne-Sophie Gosselin Grenet), as well as mouse anti ɑ-tubulin (Sigma-Aldrich T9026) antibody were incubated overnight at 4°C, followed by a 2 hours incubation with respective Horseradish peroxidase-coupled secondary antibodies (anti-mouse IgG ab97023 and anti-rabbit IgG ab97051, Abcam). Luminescence was then detected using SuperSignal West Pico PLUS Chemiluminescent Substrate (Thermo Fisher Scientific). The ImageJ software was used to perform linear contrast enhancement.

### Fluorescence microscopy

Ld652Y cells were cultured in chambered, tissue culture-treated polymer coverslips (μ-slide 4 well, Ibidi) and infected with JcDV stock at a MOI of 10. At 24 hours post infection, cell monolayers were rinsed twice with PBS, fixed in PBS-PFA (2%) for 10 min, permeabilized in PBS with triton X-100 (0.05%) for 5 min, and incubated in PBS-BSA (1%) for 30 min. Cells were then labelled with primary anti-RNAP-II antibodies (CTD Antibody, clone 8WG16, Sigma-Aldrich 05-952-I) and anti-VP (a gift from Anne-Sophie Gosselin Grenet) (26), overnight at 4°C. The next day, cells were washed with PBS-BSA (1%), and jointly labelled with Alexa fluor A488 and A568 conjugated antibodies (A110018 and A11031, Thermofisher scientific) for 2 hours at RT. Nuclei were labelled with a PBS-Hoechst solution at 2 μg/mL for 5 min at RT and samples were covered in mounting medium (ProLong P36930, Thermofisher scientific). Labelled cells were analysed with an inverted LSM 980 confocal microscope with Airyscan 8Y (Carl Zeiss Ltd.). For the co-localisation analysis, independent replicates were analysed with the Icy software (27) on the whole cells.

## RESULTS

### Accurate and cost-effective quantification of transcription errors by targeted sequencing

Accurate quantification of transcription errors necessitate deep sequencing of both RNA and DNA with high coverage to detect low frequency events. Two strategies can be considered to achieve this, non-targeted and targeted RNA sequencing. The first strategy provides information at the transcriptome scale, but requires massive sequencing throughput to reach sufficient sequencing depth, and measuring the transcription fidelity of low-expressed transcript is therefore limited. Targeted-sequencing, in contrast, provides higher sequencing depth at selected locus, and is more cost effective to assess transcription on a focused gene selection. Targeted sequencing can be achieved by enriching a sample using a set of DNA bait designed to bind the molecules of interest (28). Here we developed a more direct and economical approach that uses targeted PCR to specifically amplify loci of interest. This straightforward method is easy to implement yet poorly scalable and therefore only applicable to relatively small regions of interest.

Regardless of the approach, a major limitation in the detection of rare nucleic acid synthesis errors is that errors produced by sequencing workflows typically occur at rates that are orders of magnitude lower. Multiple technologies have been developed that eventually produce a consensus from multiple reads of the same molecule and therefore to accurately deconvolve actual mutations from sequencing errors (29, 30). To achieve a targeted version, we developed a two-step PCR procedure in which the first step uses primers containing random nucleotides to produce Unique Molecular Identifiers (UMIs) that tag the reacted single DNA or mRNA molecules. The first couple of primers also introduces half the adapter sequences required for downstream sequencing, thereby providing specific handles for the second-step PCR, which amplify the UMI-tagged molecules and introduce the rest of the sequencing adapters as well as sequencing indices required for multiplexing (**Figure 1B**). As uniquely tagged molecules are amplified and sequenced multiple times, UMI-based clustering is used to derive a consensus sequence for each molecule and unambiguously call mutations according to defined thresholds.

We applied this workflow to quantify and contrast DNA replication and mRNA transcription errors upon infection of Ld652Y insect cells by the JcDV densovirus in a set of target regions of interest (ROI) spanning both host and virus (**Figure 1A**). Briefly, Ld652Y insect cells were infected with JcDV at a high multiplicity of infection (MOI). DNA and mRNA were sequentially extracted from infected cells at 24 hours post infection in order to minimise DNA replication events. ROIs were then amplified from these samples using the two-step procedure described above and sent for multiplex sequencing. We developed a custom bioinformatic pipeline, UMIconsensus to process the resulting sequencing reads, derive UMI-based consensus reads and further characterise the nature and frequencies of true mutation (**Supplementary Figure S1 and Supplementary Table S1**). An exemplary mutational profile displaying the single nucleotide mutation frequencies derived from 1.3 x 10^7^ consensus sequences is shown in **Figure 1C**.

### Selective transcription fidelity between the host and viral genomes

We focused our analysis on 7 ROIs distributed over the two viral transcripts (representing 20% of the viral transcriptome) and 7 representative ROIs of the host transcriptome in the EF1 elongation factor, Argonaute RISC Catalytic Component 2 and Ubiquitin–selected, as well as recently annotated *Lymantria dispar* transcripts (**Supplementary Table S2**) from the study of Sparks *et al.* (31), which cover distinct cellular pathways and have different expression patterns (**Supplementary figure S2**) (32, 33). We observed a non-uniform distribution of mutations across mRNA positions, suggesting that errors are not fully random, independently of the sequencing depth (**Supplementary figure S3**). Mutations in the corresponding DNA were much less frequent and often differed in nature (**Figure 2A**), confirming that the observed RNA mutations are not mere consequences of previous DNA mutations and do reflect a distinct biological process. Globally, the different ROIs display distinct distributions of substitutions, deletions or insertions events in mRNA, but not in DNA (**Supplementary figure S4 and Supplementary figure S5**), and were reproducible across biological replicates (**Supplementary figure S6**). Most importantly, we observed a staggering 10-fold lower transcription fidelity in viral genes compared with cellular genes overall (**Figure 2B and 2C**), suggesting that about 1 in every 3 viral transcript comprise at least one mutation, and for which we verified no dependencies on viral infection (**Figure 2D**). ROI in viral genes displayed variation of mutation frequencies, even when belonging to the same mRNA transcript. For instance, we observed a 10-fold difference in average mutation frequency between region VP:763-860 (2.4×10^−4^ errors/nucleotide) and VP:1054-1140 (3.8×10^−5^ errors/nucleotide), which are located 300 nucleotides away on the same transcript. Performing this analysis at 24 hours or at 5 days post infection of insect cells reveal a similar mutational rate for all positions analysed (**Figure 2E**). Along with our data on DNA mutations, this time-dependent analysis demonstrates that mutations detected in mRNA are phenotypic mutations, arising during transcription rather than during DNA replication. To confirm the selective transcription fidelity of the viral genome, we engineered a vector bearing a complete viral genome (**Figure 2F**), with fragments of cellular genes (~160 nucleotides) inserted in place of a highly mutated ROI, VP:763-860. Because this engineering strategy alters the coding sequence of the VP gene, we also translocated a copy of the VP gene outside the viral genome, to provide a functional complementation. Such a vector allows viral gene transcription, genome replication and virus production. Downstream of the cellular gene fragment, we also introduced a 18 nucleotides priming site that allowed us to selectively sequence the cellular genes in transfected cells, either originating from the host genome or from the insertion in the viral genome. Deep sequencing of mRNA was performed at 24 hours post transfection of insect cells. Remarkably, insertion of cellular gene fragments within high mutation frequency associated-loci in viral genes decreased their transcription fidelity (**Figure 2G**). Altogether, these results demonstrate that transcription fidelity in JcDV-infected cells is a selective process, depending on the DNA template origin.

**Figure 2.**
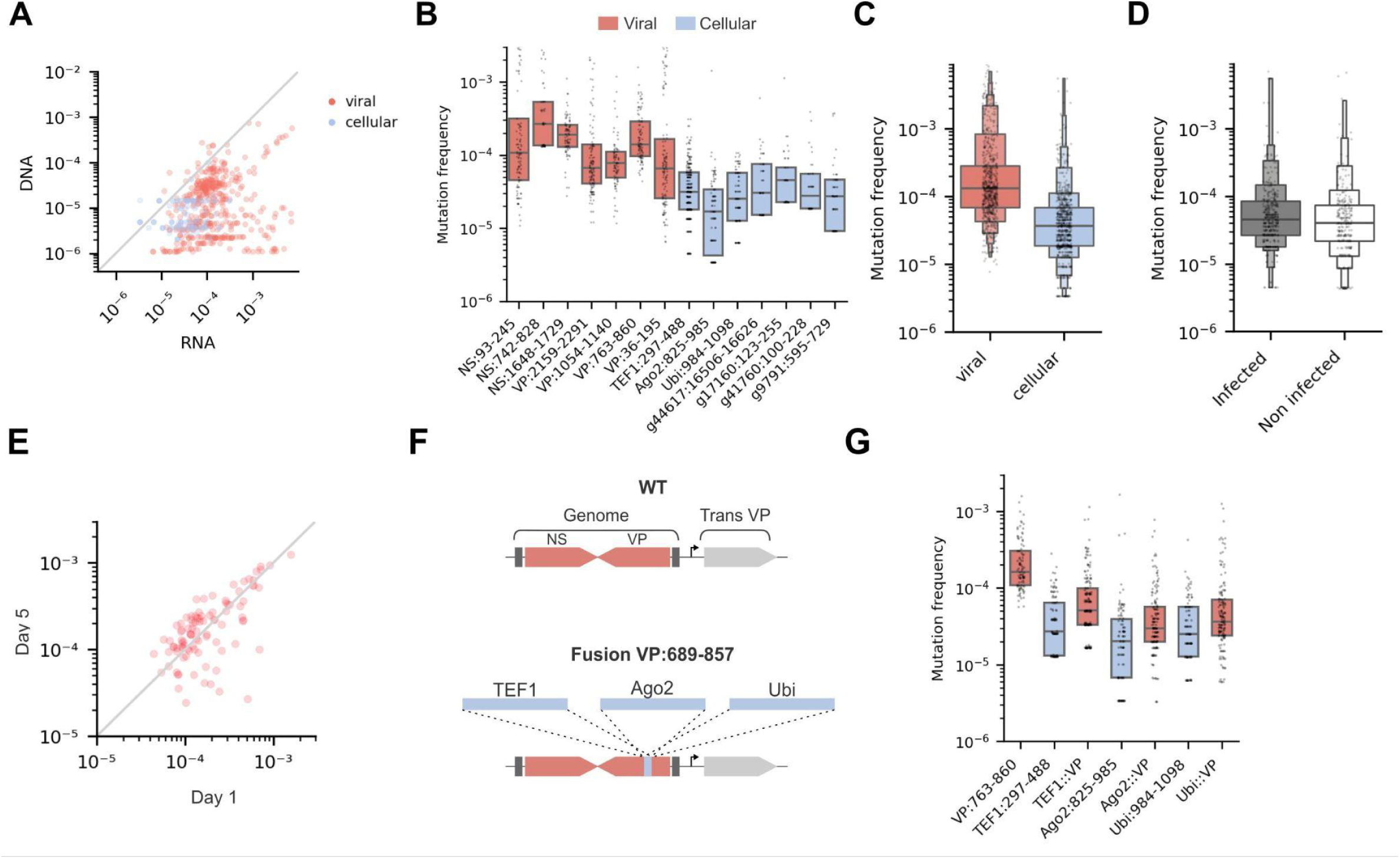
Transcription fidelity in host and viral genomes. (A) Comparison of the substitution frequencies measured in mRNA and DNA. Each point represents a position in the different ROIs, the grey line represents y=x. (B) Mean substitution frequencies across all positions within ROIs from the virus (red) and the host (blue). Vertical lines represent the 95% confidence interval (ci). (C, D) Mean substitution frequency in all targeted ROIs from (C) the viral (grey) and the host (white) genomes, or (D) from the host genome only in presence or absence of infection. Each point indicates a position. (E) Comparison of the substitution frequencies measured in mRNA ROI VP:763-860 at 1 and 5 days post infection. Each point represents a position in the ROI, the grey line represents y=x. (F) Diagram showing defective genome engineering. The JcDV genome is in red, surrounded by ITR sequences (dark grey). Inserted cellular gene fragments are in blue. A function copy of the VP gene (light grey) was inserted in trans. (G) Mean substitution frequencies across all positions within host gene fragments from the host genome (blue) or inserted in the viral genome (red). Substitution frequency measured in the viral genome in absence of exogenous insert is represented on the left (VP:763-860). Each point indicates a nucleotide position in ROIs. For (B,C,D,G), horizontal lines represent median and quartiles.

### Viral capsid proteins drive low transcription fidelity of viral genes

To determine whether the selective transcription fidelity in viral genes is a process driven by the presence of viral proteins, we endeavoured to measure the transcription fidelity in defective viral genomes. We generated three vectors containing viral genomes, each bearing one early stop codon that prevented the production of either NS3, both NS1 and NS2, or VP4, which affect all VP isoforms for the later (**Figure 3A**). Transfection of these vectors in insect cells and analysis of protein expression by western blot revealed that all constructs affected the production of capsid proteins, either by strongly decreasing VPs expression (NS1/NS2 stop) or by preventing it (NS3 stop, VP4 stop) (**Figure 3B**). Similarly, genomes with a truncated NS3 prevented the correct expression of NS1 and NS2 proteins. In contrast, the absence of VP4 (VP4 stop) did not affect the expression of NS1 and NS2 proteins. NS3 production could not be tested due to the failure of specific antibody development. We then measured the transcription fidelity for two ROIs of the VP gene in mRNA produced by defective genomes. These two ROIs were associated with a high (VP:763-860) or a low (VP:1054-1140) mutation frequency in the wildtype (WT) JcDV genome (**Figure 2B**). Analysis of mutation frequencies at each position within the two ROIs revealed that all early stop codons introduced in the viral genome induced a strong decrease of the transcription fidelity of viral genes, compared with the WT virus (**Figure 3C**). This was particularly striking for mRNAs produced by the VP4 stop construct at the VP:1054-1140 ROI, which had an error frequency of 1.9×10^−5^ mutation/nucleotide, in contrast to 8.1×10^−5^ mutation/nucleotide for the WT viral genome. We also quantified the viral mRNA and DNA levels in transfected cells using a RT-qPCR approach, and calculated the ratio of viral mRNA levels over viral DNA levels (**Figure 3D**). This showed that all constructs had similar levels of mRNA transcription. Altogether, these experiments revealed a viral origin of the selective transcription fidelity observed in viral mRNAs, which can be tuned by the expression of viral proteins.

**Figure 3.**
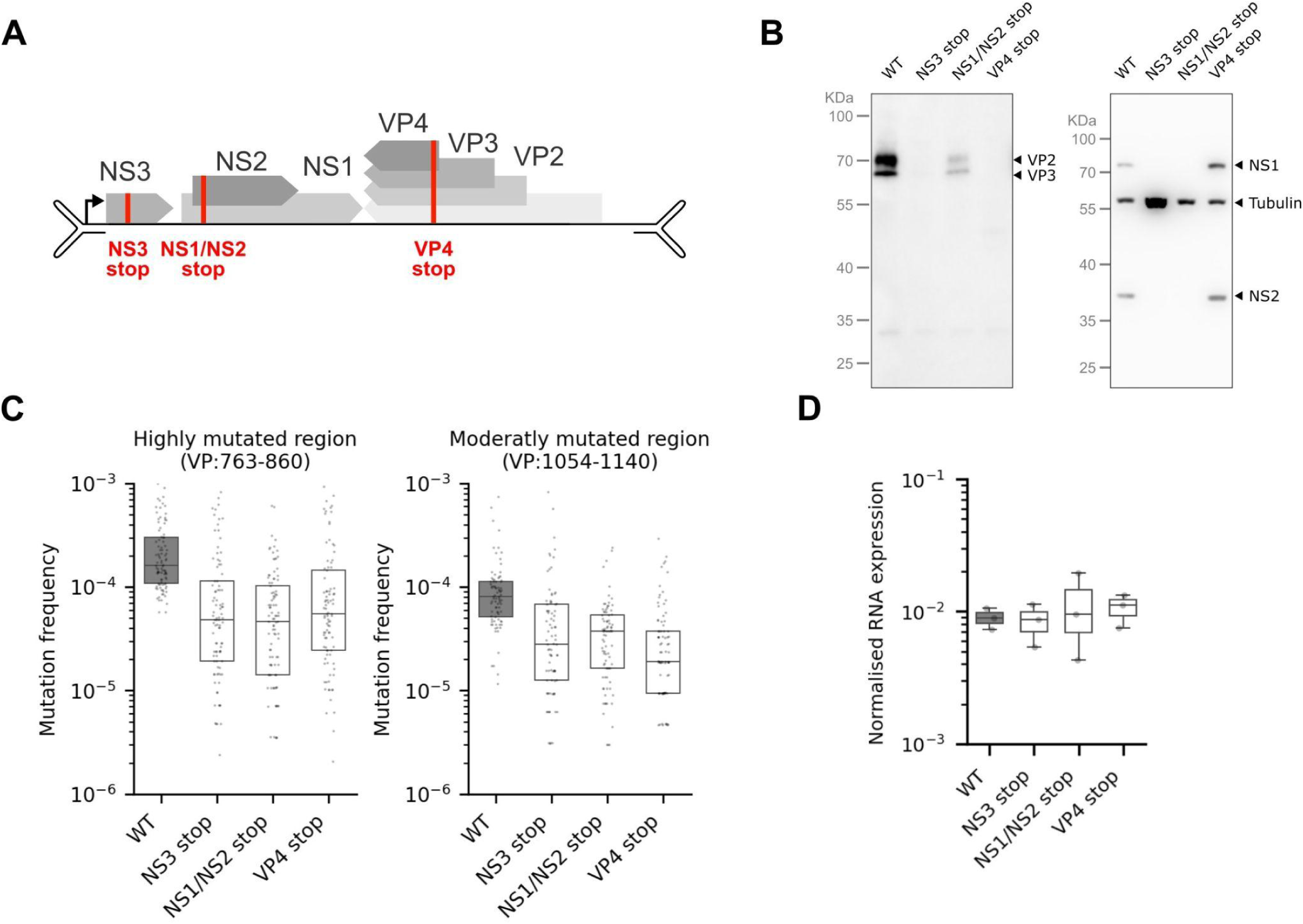
Low fidelity is induced upon viral protein expression. (A) Diagram of the JcDV genome indicating position of stop codon introductions to generate defective virus. (B) Western blot assay targeting VP isoforms (left) or non structural NS1 and NS2 proteins, and tubulin as loading control (right) in cell lysate after transfection with wild type (WT) or defective virus genomes. (C) Mean substitution/nucleotide frequencies across all positions within ROI VP:763-860 (left) or VP:1054-1140 (right) for mRNA transcribed by wild type virus (dark grey) or defective virus (white). Each point represents a nucleotide position in ROIs. (D) mRNA expression levels in cells infected with wild type (dark grey) and defective (white) virus particles. Each point indicates replicate experiments (N=3). For (C, D), horizontal lines represent median and quartiles.

### The host transcriptional machinery interact with viral capsid proteins

The host RNAP-II is responsible for the transcription of most mRNA in eukaryotic cells, and is used by the JcDV genome for transcription of viral genes as well (34, 35). Here, we observed that viral proteins are able to alter the error frequency made by the RNAP-II during viral gene transcription. We thus wondered whether viral proteins can interact with the RNAP-II in the nucleus of infected cells. To test this, we performed co-immunoprecipitation assays on nuclear proteins in cells infected with JcDV 24 hours after infection followed by western blots (**Figure 4A**). Our data show that the material coprecipitated with an antibody targeted to the RNAP-II complex contains VP, but no NS viral proteins. Conversely, co-precipitate isolated with the anti-VP antibodies contains the RNAP-II complex, while those obtained with anti-NSs antibodies do not. These results strongly suggest an interaction between the viral capsid proteins and the transcription complex. Although not well documented, VP proteins may be capable of selectively binding the viral genome for packaging. The apparent interaction between these two species could thus merely be mediated by their binding to the same viral DNA molecule. To explore this, we performed the experiment again adding a DNase I treatment prior to immunoprecipitation and obtained the same results (**Figure 4B**). To further explore this interaction between viral capsids and the RNAP-II in insect cells, we conducted an immunolabeling assay of VP and RNAP-II coupled with high resolution confocal imaging of the nucleus in infected cells at 24 hours post infection (**Figure 4C**). As characteristic of densoviruses, we observed a massive reorganisation of the chromatin within the nucleus upon JcDV infection (36). Capsid proteins are mainly localised at the periphery of the nuclear envelope, resulting in striking ring-like patterns. Consistent with the reorganisation of the chromatin, infection by JcDV dramatically alters the otherwise relatively even nuclear distribution of RNAP-II pressing it to the borders of the nucleus. As a result, we could detect a positive colocalization of fluorescence signals for RNAP-II and VP proteins, with a mean Pearson coefficient of 0.61 per cell analysed (**Figure 4D**, **Figure 4E**). Altogether, these data suggest that viral capsid proteins are able to interact with the RNAP-II complex, which provides a mechanistic basis for the impact of JcDV on its own transcription fidelity.

**Figure 4.**
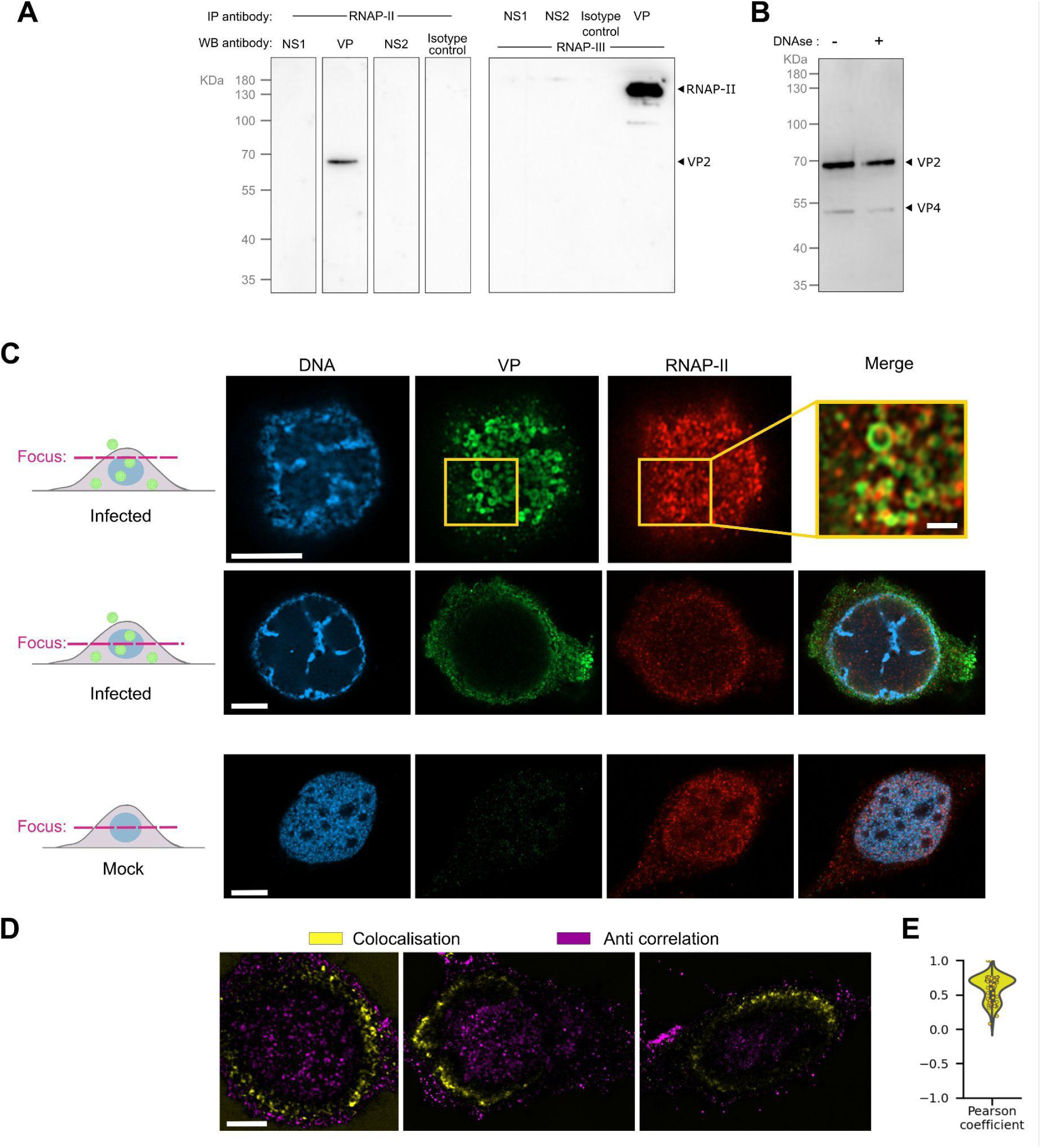
Interaction between viral proteins and the host transcription machinery. (A) Immunoprecipitation of RNAP-II (left) or viral proteins (right) from infected cell lysates using specific antibodies or sera, and subsequent detection of viral proteins (left) or RNAP-II (right) in precipitates by western blot assay. (B) Immunoprecipitation of RNAP-II after DNase treatment of the cell lysate, and detection of VP protein by western blot assay in the precipitate.(C) Subcellular localisation of VP (green) and RNAP-II (red) in infected insect cells (24 hours post infection) observed by super resolution confocal microscopy focused at the top (top row) or centre (middle row) of the nucleus. Non-infected cells imaging were also imaged (bottom row), as well as chromatin labelling (Blue). White scale bars indicate 5 µm, except in the image magnification (yellow square) where it indicates 1 µm. (D) Coloured representation of colocalization (yellow) or anti-colocalization (magenta) between RNAP-II and VP images. (E) Distribution of the mean Pearson correlation coefficient between VP and RNAP-II in infected cells (N=61).

### Sequence composition finely modulate the transcription fidelity

Next, we also analysed the frequency of each mutation type (**Figure 5A**). We observed more frequent transitions (A to G: 9.9×10^−4^ errors/nucleotide and G to A: 7.2×10^−4^ errors/nucleotide for most frequent nucleic acid transitions) than transversions (C to A: 2.5×10^−4^ errors/nucleotide, C to G: 1.2×10^−4^ errors/nucleotide for most frequent nucleic acid transversions). Insertions and deletions events were less observed compared with substitutions. In addition, we observed a large heterogeneity in distributions among the mutation types across target ROIs (**Figure 5B**). Some mutations are always detected at high frequency, regardless of their exact position, such as G to A transitions. In contrast, some mutations, such as A to G, displayed a multimodal distribution with positions associated with lower or higher mutation frequencies. This suggests that, in addition to the role of capsid proteins in tuning the transcription error frequency, the local context within the template gene could also be a factor influencing the error frequency. Among other factors, we observed that the presence of long homopolymers, above 5 nucleotides, within the gene decreases transcription fidelity (**Figure 5C**). Similarly, the A+T content was also correlated with a decrease in transcription fidelity, when comparing the A+T content for all 9 nucleotides long windows, with their respective mutation frequency (**Figure 5D**). To decipher more precisely the impact of the nucleotide context upstream and downstream each position, we analysed all nucleotides 3 mers that were present within our targeted ROIs (**Figure 5E**). We compared their median mutation frequency (**Figure 5F**), and observed a wide distribution of mutation frequency associated. Some 3 mers were systematically associated with high transcription fidelity, especially those with a middle adenosine (AAG, TAA, TAC) while others were associated with poor sequence conservation (CTG, TTG, CTG). These observations indicate that the sequence context is a potential fine regulator of the local transcription fidelity in hypermutated viral transcripts.

**Figure 5.**
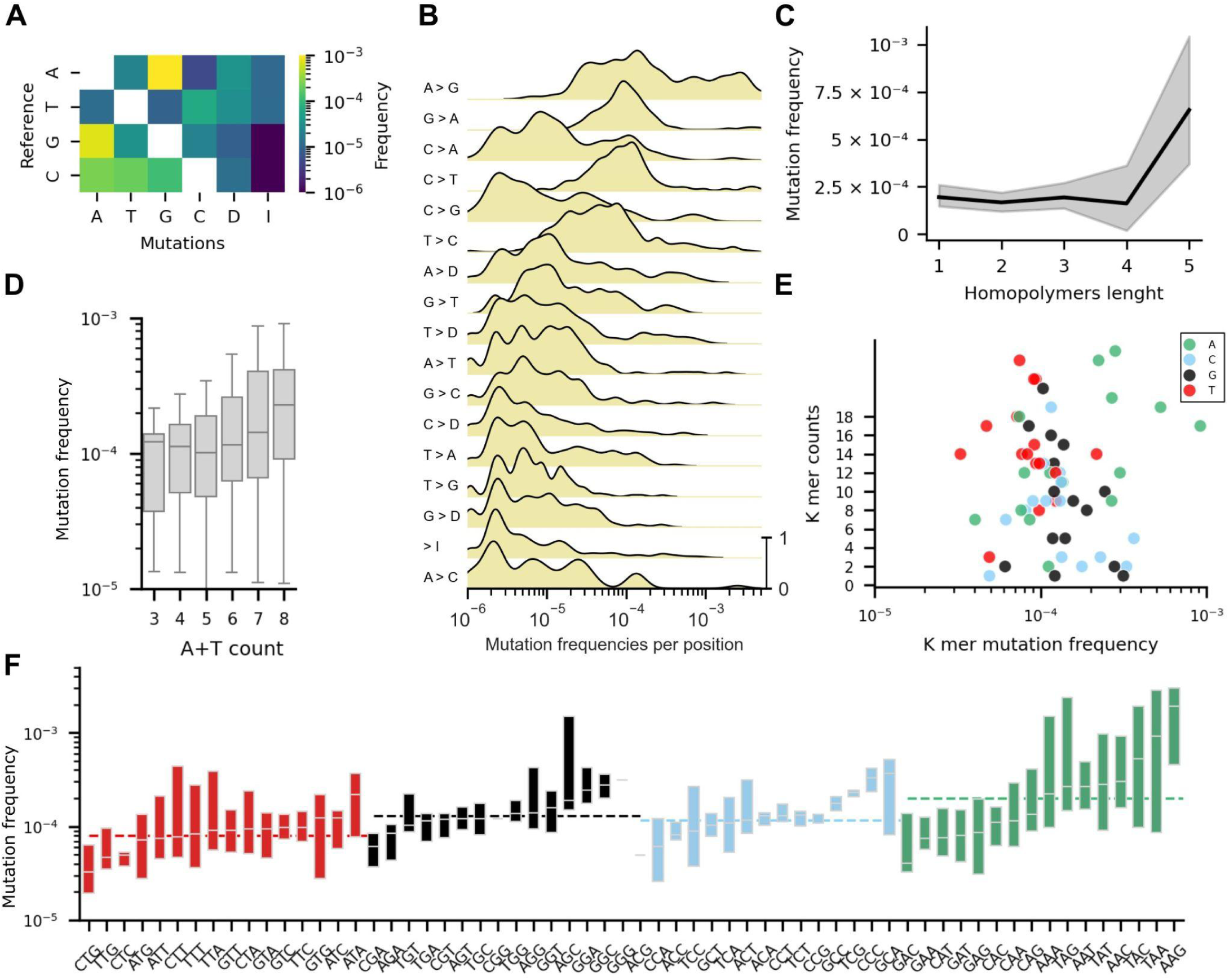
Role of the sequence context on shaping the virus transcription fidelity. (A) Frequency of mutations/nucleotide for each mutation type. D: Deletion, I: Insertions. (B) kernel density estimate of mutations/nucleotide frequencies per position in the viral genome. Each row shows the distribution of a specific mutation type (indicated on the left). Y-axis scale is indicated at the bottom right. (C) Evolution of the mutation/nucleotide frequency based on the homopolymer length. Dark line represents the mean frequency, and the 95% confidence interval is plotted in grey. (D) Evolution of the mutation/nucleotide substitution based on the A and T nucleotides content in each 9-nucleotides window of the viral ROIs. (E) Relation between occurrences for each 3-mer nucleotide sequence detected in viral ROIs with their median frequency of mutation. (F) Distribution of mutations/nucleotides observed for each 3-mer. The middle nucleotide only is used as a reference to measure the mutation frequency. Dashed horizontal lines represent median mutation frequency for each pool of kmer based on their middle nucleotide. For (D, F), horizontal lines in each box represent median and quartiles.

### Impact of low transcription fidelity on protein translation

After investigating the causes of low transcription fidelity in viral genes, we shifted focus to its effect on protein translation. The consequences of a single nucleotide mutation is strongly dependent on the topology of the genetic code. Using the mutation frequency measured at each nucleotide position within the ROIs (**Figure 1C, Supplementary figure S4**), we deduced the frequency of codon changes. Our data indicate that 42% of mRNA errors translate to non-synonymous changes (**Figure 6A**), with a median mutation frequency of 4.4×10^−5^ and 3.2×10^−5^, for synonymous changes and non-synonymous changes respectively. For the JcDV protein which comprise seven proteins (NS1, NS2, NS3 and VP1, VP2, VP3, VP4) with 3323 amino acids, this equates to a mean of 7.4% (7.4 ×10^−2^ mean mutation/protein) of translated proteins harbouring a transcriptional mutation. We then predicted the amino acids substitution matrix (**Figure 6B**). For this, we took into account only amino acid changes requiring one substitution, since a majority of mutations observed were single mutations within the targeted ROIs. The most mutated amino-acid observed was the valine (6.7×10^−3^ mutations/amino acid), lysine (3.9×10^−3^ mutations/amino acid) and threonine (3.1×10^−3^ mutations/amino acid). Valine codon mutations were predominantly mutated toward isoleucine codons, relying on the nucleic acid transition of the first codon base (G to A).

**Figure 6.**
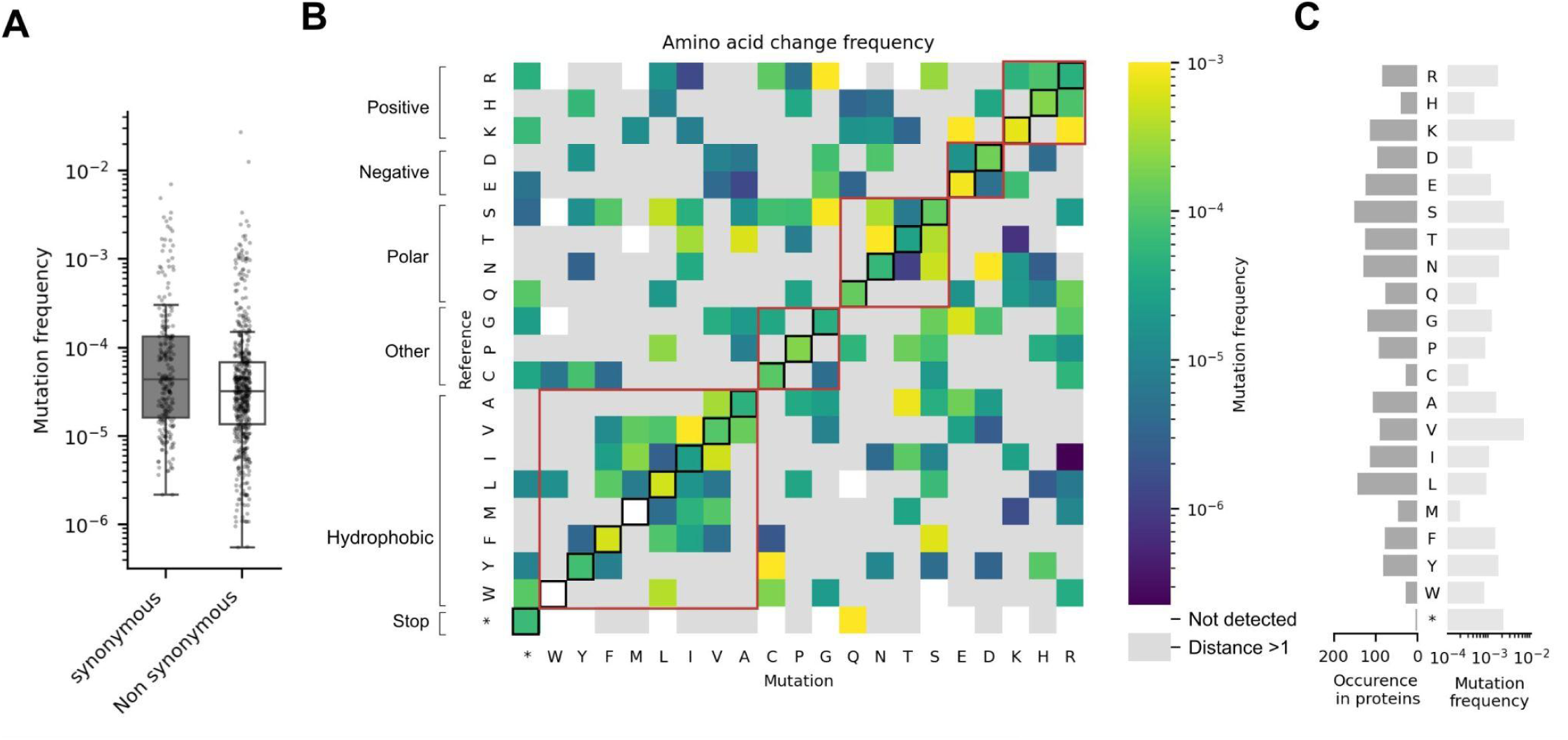
Impact of low transcription fidelity on viral proteins. (A) Frequency of synonymous and non-synonymous mutations/codon observed in viral ROIs. Each point represents a codon. Horizontal lines represent median and quartiles, while whiskers extend to the rest of the distribution. (B) Heatmap showing the change/amino-acid frequency for each amino acid detected in viral ROIs. Rows indicate reference amino acids and columns indicate amino acids changes. Dark edge squares represent synonymous mutations, while red edge squares groups amino-acids sharing similar side chain properties (indicated on the left). Only amino acid change requiring single nucleotide substitutions are analysed (otherwise coloured in pink). Amino acid changes not detected in our data are coloured in white. (C) Comparison of amino-acids occurrences (left) in viral proteins, compared with their respective mutation frequency (right).

Similarly, the next most observed amino acid changes were threonine to asparagine (C to A transversion), lysine to arginine (A to G transition) and lysine to glutamic acid (A to G transition). This is in line with the preferential substitution of A to G, G to A, and C to A observed (**Figure 5A**) in viral transcripts, and illustrates how factors driving transcription fidelity induce biassed amino-changes in proteins. We did not observe a strong correlation between amino acid occurrence in viral proteins and their respective mutation rate (**Figure 6C**). For instance, aspartic acid, leucine and glutamine amino acids were poorly mutated compared with their occurrences in the JcDV genome. This observation is striking for the leucine amino acid, which is encoded by more codons (6 codons), compared to the others (2 codons), providing more putative exploration of the amino acid sequence space. Similarly, the codon stop reversion of the NS3 genome toward a glutamine was among the top mutations recorded, whereas we only recorded one codon stop in ROIs (NS:742-828). Such mutant extends the NS3 coding frame, and induces production of a NS3:NS1 fusion protein, because only 6 nucleotides are separating the two coding sequences. Overall, our analysis underscores the profound impact of transcriptional errors on protein synthesis, and indicates tight regulation of amino-acid changes despite the high level of transcription errors.

## DISCUSSION

Most mutations are neutral or deleterious for the fitness of an organism, and even in the case of beneficial mutations, their likelihood of appearance is independent of their phenotypic consequences. However, accumulating evidence suggests that organisms can increase their mutation rate in response to environmental stress (37, 38), but at the genome scale rather than on specific genes or loci. Established for heritable mutations, this paradigm may not necessarily apply to phenotypic mutations, which can exhibit variable mutation rates within a genome as our findings in JcDV transcription suggest. These corroborate observations made in yeast and human cells, where the fidelity of transcription is different within the transcriptome, but also between regions in a transcript (5, 6). Our investigation extends this concept by demonstrating that the transcription error rates can be selectively tuned by a parasitic genetic entity such as a virus, independent of the host genome despite both genomes rely on the same transcription machinery.

To measure mutations arising during transcription, we derived robust and high quality consensus sequences of unique mRNA molecules isolated from infected cells. For this, we developed a targeted high-throughput sequencing strategy based on the clustering of Unique Molecular Identifiers (UMI) (29) associated with stringent analytical criteria, such as imposing high quality score thresholds and elevated number of reads per UMI cluster (**Supplementary Figure S1**). This approach enables a reliable and cost-efficient analysis by focusing the sequencing throughput on regions of interest. Unlike some non targeted strategies, such as the CircSeq method (30), it does not account for errors occurring during the reverse transcription of mRNAs. Such errors should impact all samples similarly and would only question the absolute nature of the numbers reported here. As the reported error rate for reverse transcriptase are orders of magnitude lower than those we observed (6.3 ±1.2 x 10^−5^ error/nucleotide for the Moloney Murine Leukemia Virus Reverse Transcriptase) (39–41), such reverse transcription error can be safely neglected.

Our analysis revealed that the error rate of transcription is on average 10-fold higher in viral as compared to host genes (**Figure 2C**), while the transcription fidelity of host genes is not impacted by infection (**Figure 2D**). While this discrepancy could be argued to stem from differential intrinsic propensities of the investigated sequences to elicit transcriptional errors, we show that the very same host genes sequences experience increased errors when transcribed from within the viral genome (**Figure 2G**). This unambiguously demonstrates that the viral genome is selectively subject to an hypermutator phenotype at the transcription level. Consistent with previous findings (6), our study did not reveal a direct relationship between mutation frequency and gene expression levels.

Suppressing the expression of individual viral proteins systematically resulted in a fidelity increase (**Figure 3C**), supporting a mechanistic link between the virus and the mutator phenotype. All of these manipulations are expected to compromise viral replication. A lack of replication, however, cannot explain these results since the mutations observed are *bona fide* transcription errors, which are not associated in quality or quantity with replication errors that could be observed in the viral genome. Another common trait of these manipulations is the sharp reduction in VP expression level, which could mediate the observed effect (**Figure 3C**). Our prior work established that truncation of capsid proteins halts viral genome replication without affecting transcription (42).

Reciprocal co-immunoprecipitation and colocalization experiments further strengthened the role of VP proteins by supporting its interaction with the host RNAP-II complex. Identification of interactions between the capsid protein and RNAP-II sheds light on the mechanism by which capsid proteins may modulate mutation rates. Using super-resolution microscopy, we detected the spatial arrangement of VP in ring-like formations at the nuclear periphery (**Figure 4C**), suggesting that VPs may sequester RNAP-II within these structures. This spatial segregation could underpin a differential transcription error rate between host and virus, filtering the genomic mutation landscape. Moreover, RNAP-II and VP interaction can be indirect. Other proteins are known to destabilise RNAP–II, such as the TFIIS co-factor, that increase RNAP-II proofreading abilities (43, 44). The capsid protein isoforms, including the most prevalent and shortest VP4, are produced from a single gene using alternative initiation codons. Our VP targeting antibody has a strong affinity for the VP2 isoform, rendering the detection of other VP isoforms challenging, and dependent on the protein extraction method. However, in our immunoprecipitation assay,t VP4 interaction with the RNAP-II complex was confirmed (**Figure 4A**). This suggests that the VP direct interactor within the RNAP-II complex, is able to bind all VP within their VP4 region.

In our investigation of viral transcripts hypermutations, we noted a variation of transcription fidelity across transcript loci. This observation implies that factors beyond capsid protein presence are influencing the mutation rate. To explore this, we examined the sequence context of mutations. While C to T substitutions are predominant in RNAP-II mediated transcription in yeast and human cells (5, 6), we observed that G to A and A to G substitutions are favoured during viral genome transcription (**Figure 5A**). For most substitution types, such as A to G or C to A, we observed a multimodal distribution of frequency. This reveals that nucleotides within the viral genome do not always mutate with the same frequency (**Figure 5B**), suggesting a local adjustment of the transcription fidelity based on the sequence contexts. In line with this, we identified specific nucleotide 3-mers, such as AAG, TAA, TAC, that are more susceptible to mutation during transcription (**Figure 5F**), highlighting the intricate role of sequence context in determining transcriptional error rates. Local AT content is another example of transcription fidelity adjustment, possibly linked to DNA-RNA hybrid interaction or DNA secondary structures. Indeed, we found a nearly linear correlation between the local AT content and the transcription fidelity (**Figure 5D**).

In this study, we acknowledge certain limitations inherent to our methodological approach. Specifically, our targeted sequencing approach does not allow us to study the frequency of multiple mutations on distant loci, which exclude the identification of potential linkage between mutation hotspots. Nonetheless, we found that multiple mutational events within the same targeted region were rare events compared with single mutations (**Supplementary Figure S7**). Previous studies have reported a relationship between DNA methylation and transcription fidelity (5). This was interpreted as a consequence of the DNA methylation effect on expression levels (45). In our analysis, we did not find a correlation between transcription activity and fidelity (**Figure 2B and Supplementary Figure S7**). Despite this, it would remain compelling to study chemical modifications of the viral genomes and their impact on transcription fidelity. This could provide deeper insights into the mechanisms governing transcriptional accuracy and the potential role of epigenetic modifications in this process.

Phenotypic mutations, arising from errors in transcription or translation, are known to significantly impact protein folding, stability, and function, as evidenced in previous studies (4, 9, 46, 47). In our analysis of viral transcripts, a substantial proportion of mutations were found to be non-synonymous (**Figure 6A**), leading to the production of multiple variants with potentially altered functions (**Figure 6B**). This mechanism results in viral populations with genetically identical genomes, yet a fraction of individuals will exhibit phenotypic variation due to the presence of mutated proteins. For instance, we observed that mutation of the NS3 stop codon toward a glutamine was among our top non synonymous mutations recorded. This mutation enables leaky termination and extension of the NS3 protein, producing a NS3:NS1 fusion protein, since only 6 nucleotides separate the NS3 and NS1 genes. Considering our average non-synonymous mutation rate of 3.2×10^−5^ with the 3323 amino acids encoded across the JcDV proteome, we can roughly estimate that 7% of the viral proteins synthesised comprise at least one amino acid change. Also, we did not observe an association between amino acid occurrence in viral proteins and their respective mutation rate (**Figure 6C**). This reinforces our suspicion that variation of the transcription fidelity is selected by favouring mutation of specific amino acids, regardless of their frequency in the viral proteome. JcDV has a single stranded DNA genome, and mutations occurring during transcription cannot be heritable. Hence, viral capsids bearing phenotypic mutations carry and propagate a non mutated genome, raising the question whether there is an advantage conferred by phenotypic mutations. While most mutations are expected to be neutral or deleterious to the virus fitness (7, 48).

Non-heritable mutation can be useful momentarily but is not effective for selective pressure that is maintained for a longer period of time. But increased short term survival also increases the chances an heritable mutation arises and is selected. Indeed, some rare variants may occasionally confer a transient advantage over their wildtype counterparts and can be advantageous for the virus during its infection cycle, particularly after the release in environments that demand high persistence (49, 50), and across diverse physiological barriers encountered during the host infection (51). In an experiment with an error prone T7 RNA polymerase it was shown that eliciting transcriptional mutation in an antibiotic resistance gene favoured the evolution of heritable antibiotic resistance phenotype (9). Extrapolating on these findings, it is tempting to speculate that viruses could evolve phenotypic plasticity -via e.g. alteration of the host transcription’s fidelity- to provide increased chances to overcome transient roadblock to infection, improving opportunities for evolution without incurring the cost of a high mutation load.

## AUTHOR CONTRIBUTIONS

Thomas Labadie: Conceptualization, Formal analysis, Methodology, Visualization, Writing—original draft.

Guillaume Cambray: Conceptualization, Formal analysis, Methodology, Visualization, Writing—review & editing.

## ACKNOWLEDGEMENTS

We are grateful to A. S. Gosselin-Grenet for useful discussions, and for providing antibodies targeting *Junonia Coenia* Densovirus proteins.

## FUNDING

This work was funded by an ATIP-Avenir grant from CNRS to G.C.

## CONFLICT OF INTEREST

We declare no conflict of interest

## SUPPLEMENTARY MATERIAL

**Supplementary figure S1. Effect of experimental and computational parameters on the measure of transcription fidelity.** (A) Comparison of the median mutation frequency observed with either the PCR cycles to amplify the NGS library, the raw reads number obtained (input), the reads number after pair-reads merging (merged), the reads number after alignment filtering (aligned), the number of unique UMIs obtained, the number of UMI clusters obtained (using adjacency clustering method) and the average depth (read number) per UMI cluster. Each dot indicates a different NGS library. Horizontal dashed grey line indicate the threshold of mutation detection, below which the mutation rate could not be calculated (B) Influence of the minimum pair reads number (depth threshold) allowed to support a consensus sequence, and the minimum mean base calling quality score (quality threshold) allowed per consensus sequence at all positions, on the final average sequencing depth per position in ROIs. (C) Influence of the UMI clustering method (Unique = No Clustering, Adjacency, Clustering or directional) on the frequency of substitutions/nucleotide measured at all positions in mRNA (top) for the ROI TEF1:297-488 (left) or VP:763-860 (right). Reference nucleotides are indicated at each position. N indicates the number of UMI clusters (=consensus sequence) obtained with each clustering method.

**Supplementary figure S2.** Host gene expression levels in infected (grey) and non-infected (white) cells. Each point indicates quantification replicates (N=3). Horizontal lines represent median and quartiles.

**Supplementary figure S3.** Comparison of mutation frequency and sequencing depth. Frequency of nucleotides substitution measured in mRNA (top) and number of consensus sequences (bottom) at all positions for viral (VP:763-860, VP:1054-1140, NS:1649-1729) and cellular (TEF1:297-488, Ubiquitin:984-1098) ROIs. Reference nucleotides are indicated at each position (middle).

**Supplementary figure S4.** Comparison of substitution in RNA and DNA. Frequency of nucleotides substitution measured at all positions in mRNA (top) or DNA (bottom) for viral (VP:763-860, VP:1054-1140, NS:1649-1729) and cellular (TEF1:297-488, Ubiquitin:984-1098) ROIs. Reference nucleotides are indicated at each position (middle).

**Supplementary figure S5.** Comparison of deletions/insertions in RNA and DNA.Frequency of nucleotides deletion (purple) and insertion (yellow) measured at all positions in mRNA (top) or DNA (bottom) for viral (VP:763-860, VP:1054-1140, NS:1649-1729) and cellular (TEF1:297-488, Ubiquitin:984-1098) ROIs. Reference nucleotides are indicated at each position (middle).

**Supplementary figure S6. Replicated measures of transcription fidelity.** Comparison of the mutation frequencies measured in viral (VP:763-860) and cellular (TEF1, Ubiquitin) ROIs for different biological replicates. Each point represents a position in the ROI, the grey line represents y=x.

**Supplementary figure S7. Percentage of consensus sequences with N mutations.** Percentage of consensus sequences derived from each UMI cluster bearing 0, 1, 2 or more substitutions (top), deletion (middle) or insertion (bottom), for consensus derived from RNA sequencing (left) or DNA sequencing (right) and viral ROIs (blue) or cellular ROIs (orange). Horizontal lines represent the 95% confidence interval (ci).

**Supplementary Table S1. Effect of different clustering methods on UMI consensus obtained.** The initial number of reads before processing (input), after pair-read merging using NGmerge (Merged), after UMI extraction using CutAdapt (Cut), after quality filtering of UMIs (Umied), and after alignment to a reference sequence (Aligned) are detailed for four ROIs. “UMIs unique” indicate the total number of unique UMIs found before UMIs clustering. The number of UMI clusters using adjacency, cluster or directional method or without clustering (unique) are also indicated, as well as the average read numbers per UMI cluster obtained

**Supplementary Table S2.** Results of a blast analysis for LD652Y transcripts analysed in this study that are not annotated in genbank (NCBI).

## Notes

### Competing Interest Statement

The authors have declared no competing interest.

