## Supplementary_Data for "Selective manipulation of the host RNA-polymerase transcription fidelity increases phenotypic mutations in an insect *Parvovirus*": Supplementary_tables.pdf

| Algorithm steps | VP:1054-1140 | Ubi:984-1098 | VP:763-860 | TEF1:297-488 |
|---|---|---|---|---|
| Input reads (raw data) | 6890767.0 | 7491812.0 | 7574500 | 9379008 |
| Merged reads | 6710477.0 | 7442042.0 | 5610991 | 9334683 |
| Cut reads | 5780868.0 | 7435619.0 | 5442419 | 9314857 |
| Umied reads | 5731080.0 | 7407601.0 | 5388725 | 9237236 |
| Aligned reads | 5712569.0 | 7407073.0 | 5380588 | 9234872 |
| UMIs unique | 4196299 | 437447 | 1342130 | 309505 |
| UMIs clusters (*clustering*) | 4000940.0 | 156111.0 | 1306551 | 51572.0 |
| UMIs clusters (*adjacency*) | 4017904 | 268179 | 1307783 | 130259 |
| UMIs clusters (*directional*) | 4011063 | 250665 | 1309142 | 102347 |
| Average depth (*clustering*) | 349631.6 | 105827.8 | 522430.2 | 23848.1 |
| Average depth (*adjacency*) | 345047.3 | 189098.3 | 522686.4 | 77338.7 |
| Average depth (*directional*) | 345092.5 | 193724.7 | 523586.6 | 70817.0 |
| Average depth (*unique*) | 286581.3 | 238215.1 | 521692.7 | 130532.0 |

### Supplementary Table S1

| Gene ID | Description | Scientific name | Total score | Query cover | E value | Perc. identity | Acc. Len | Accession |
|---|---|---|---|---|---|---|---|---|
| g44617 | Muscle-specific protein 300 kDa isoform X6 | *Spodoptera frugiperda* | 13489 | 99% | 0.0 | 90.41% | 16968 | XP_050561727.1 |
| g17160 | Bicaudal D-related protein homolog | *Spodoptera litura* | 1045 | 100% | 0.0 | 91.17% | 589 | XP_022831601.1 |
| g41760 | Monocarboxylate transporter 12 | *Spodoptera frugiperda* | 873 | 98% | 0.0 | 78.93% | 545 | XP_035446775.1 |
| g9791 | Eukaryotic translation initiation factor 5B-like | *Spodoptera litura* | 304 | 95% | 3e-98 | 78.30% | 332 | XP_022829752.1 |

Supplementary Table S2
