## Supplementary figures and images for "Selective manipulation of the host RNA-polymerase transcription fidelity increases phenotypic mutations in an insect *Parvovirus*"

### figS1.jpg

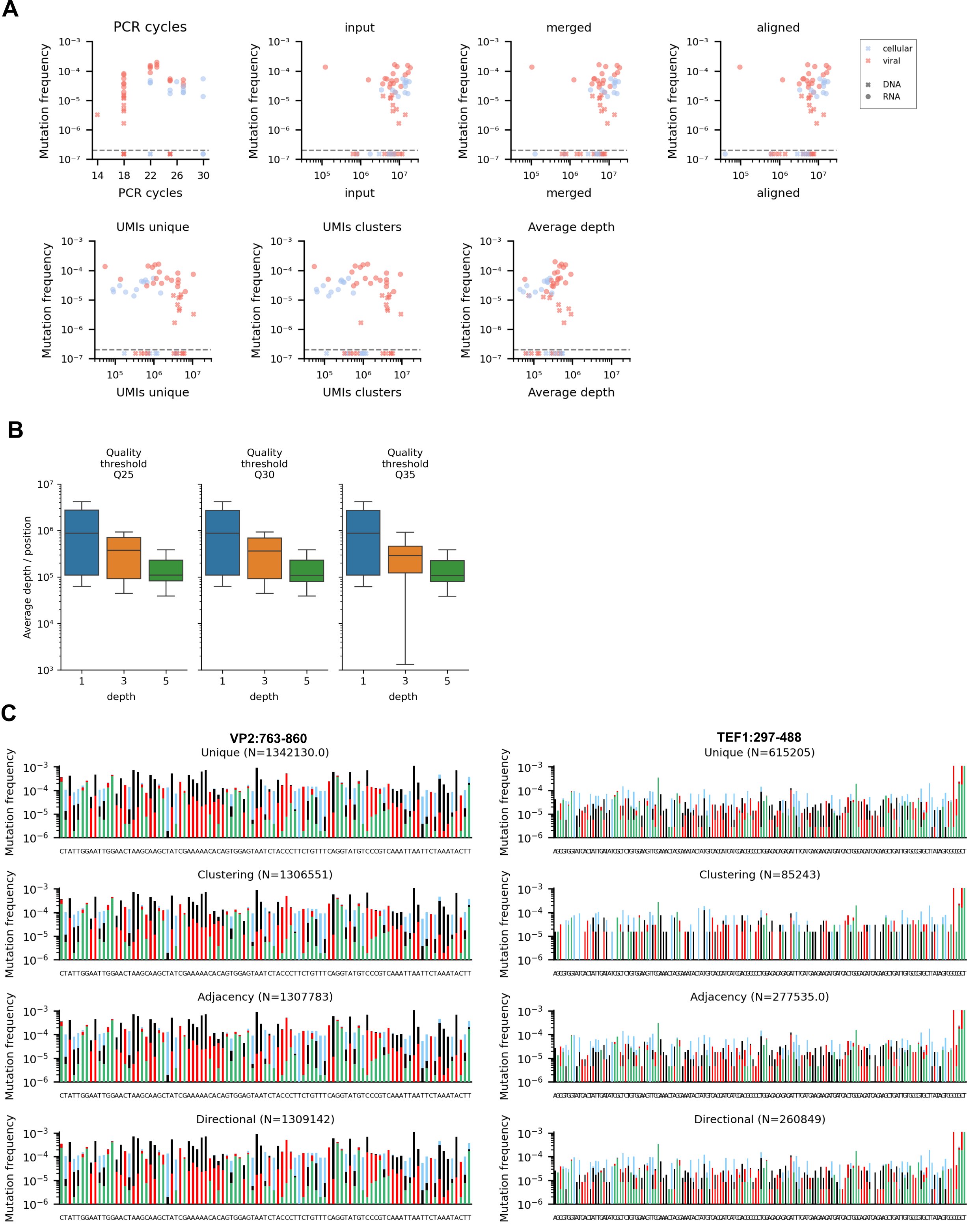

### figS2.jpg

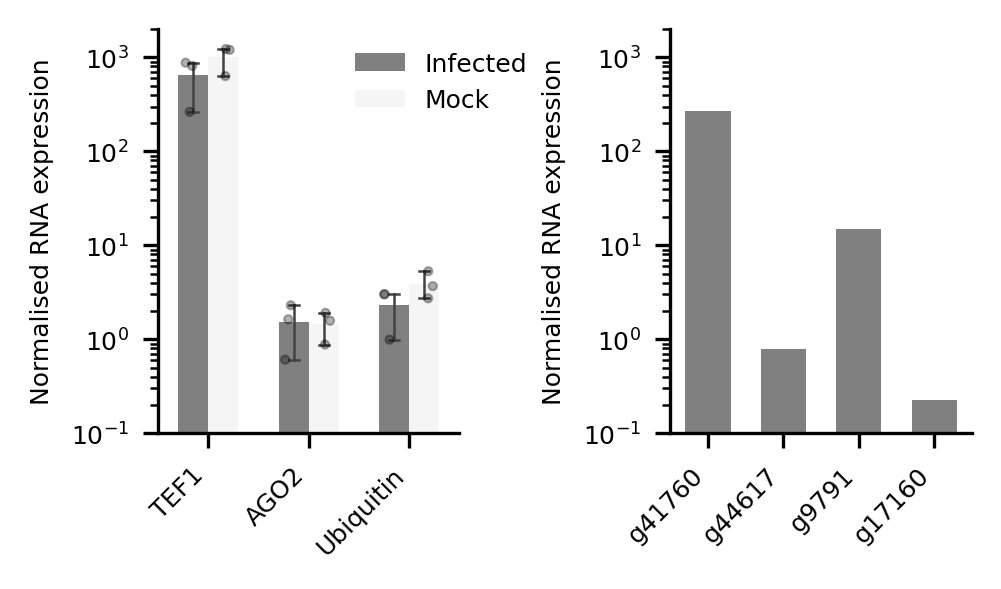

### figS3.jpg

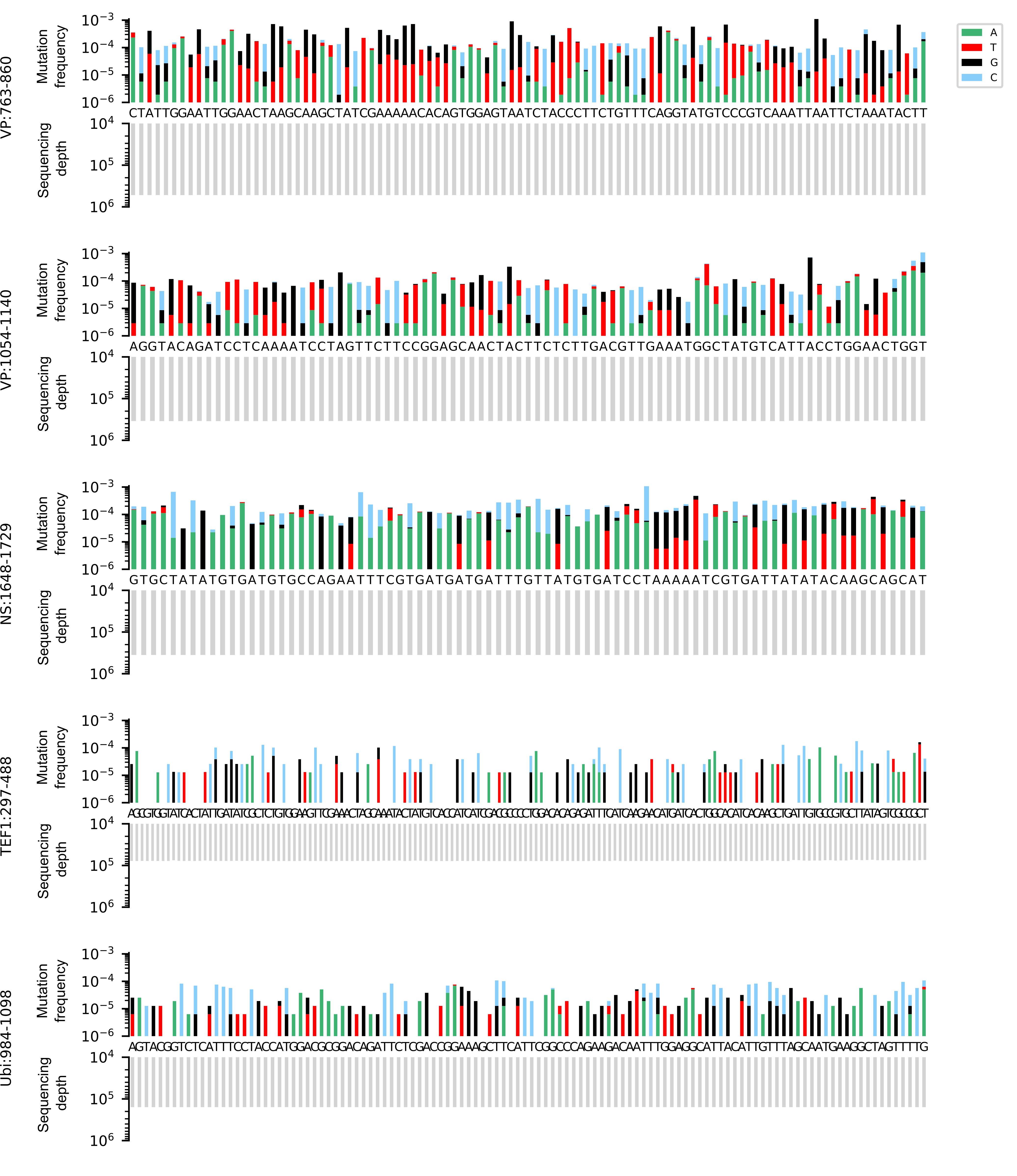

### figS4.jpg

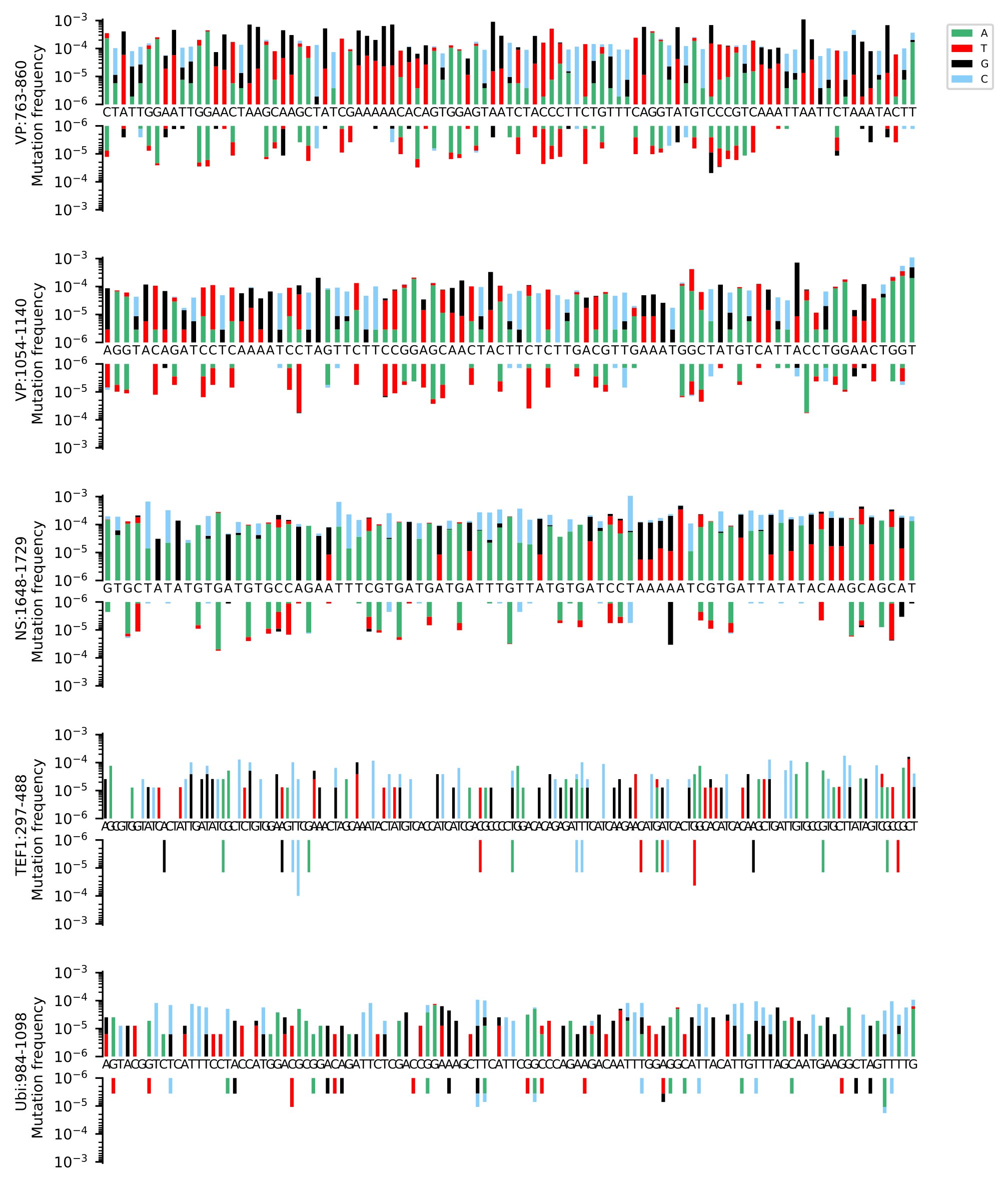

### figS5.jpg

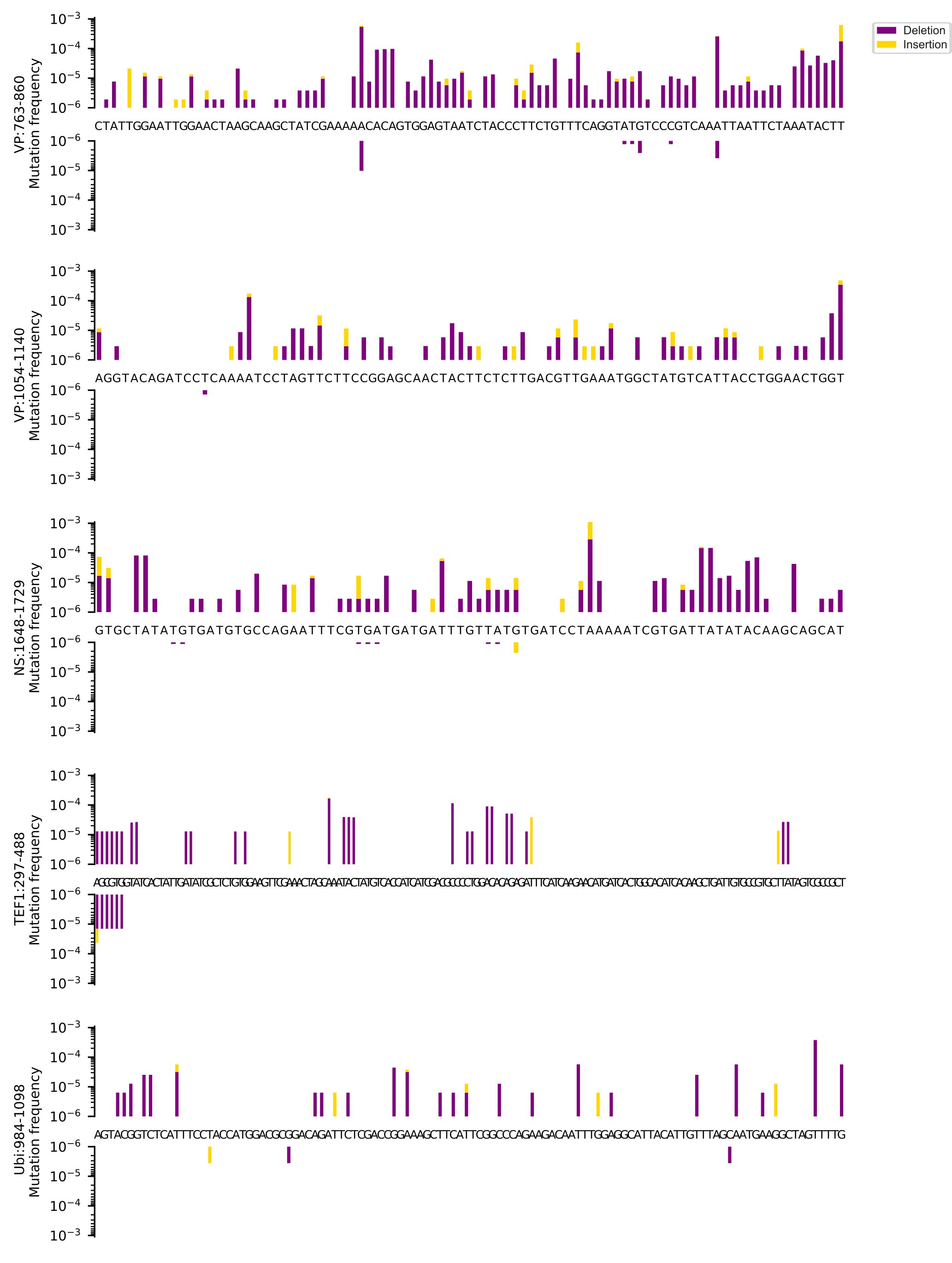

### figS6.jpg

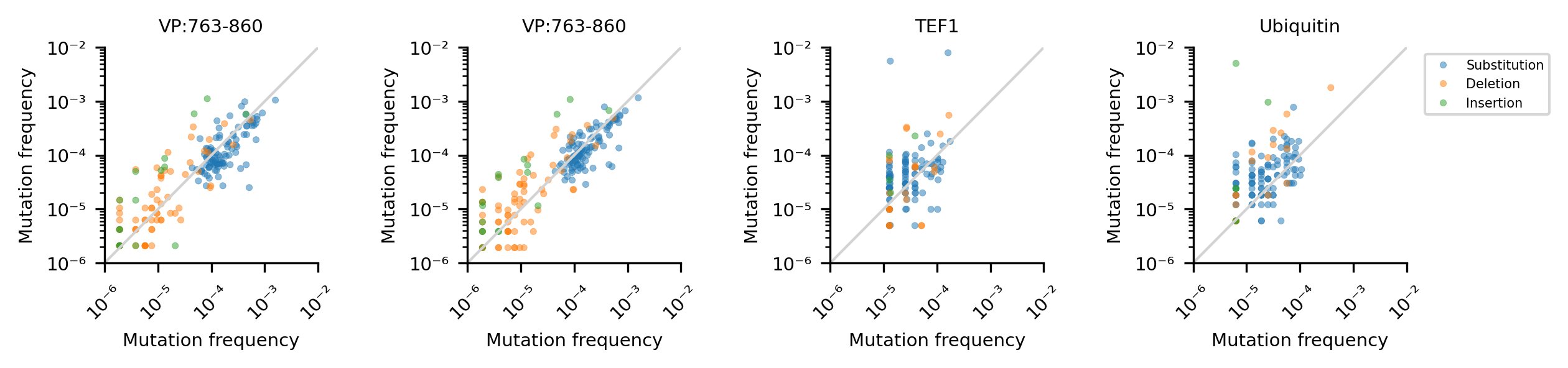

### figS7.jpg

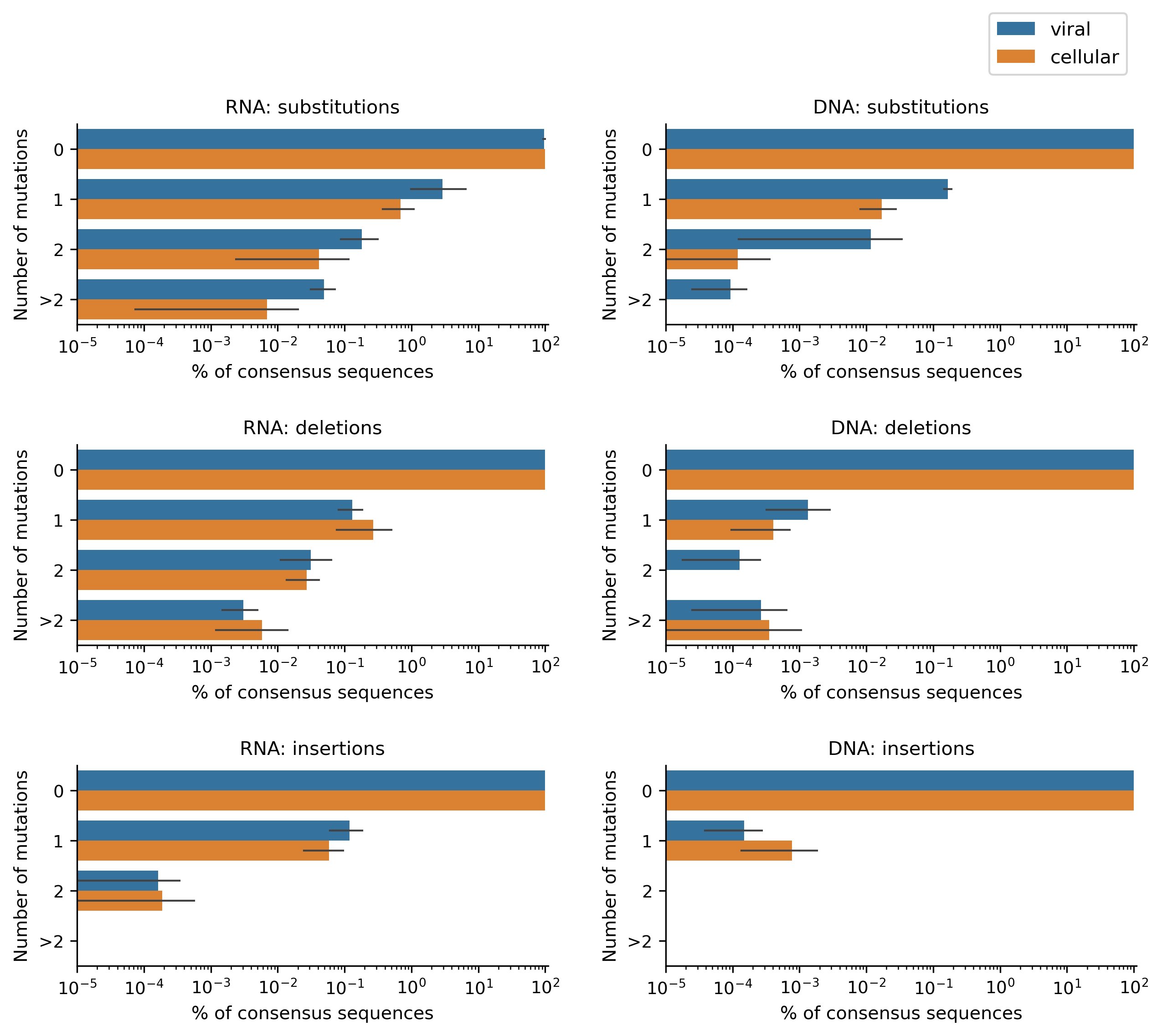
